# Invasive ant learning is not affected by seven potential neuroactive chemicals

**DOI:** 10.1101/2022.11.01.514620

**Authors:** Henrique Galante, Tomer J. Czaczkes

## Abstract

Nectar-feeding insects are often the victims of psychoactive manipulation, with plants lacing their nectar with secondary metabolites such as alkaloids and non-protein amino acids which often boost learning, foraging, or recruitment. However, the effect of neuroactive chemicals has seldomly been explored in ants. Argentine ants (*Linepithema humile*) are one of the most damaging invasive alien species worldwide. Enhancing or disrupting cognitive abilities, such as learning, has the potential to improve management efforts, for example by increasing preference for a bait, or improving ants’ ability to learn its characteristics or location. Here, we test the effects of seven potential neuroactive chemicals - two alkaloids: caffeine and nicotine; two biogenic amines: dopamine and octopamine, and three non-protein amino acids: β-alanine, GABA and taurine - on the cognitive abilities of invasive *L. humile* using bifurcation mazes. Our results confirm that these ants are strong associative learners, requiring as little as one experience to develop an association. However, we show no short-term effect of any of the chemicals tested on spatial learning, and in addition no effect of caffeine on short-term olfactory learning. This lack of effect is surprising, given the extensive reports of the tested chemicals affecting learning and foraging in bees. This mismatch could be due to the heavy bias towards bees in the literature, a positive result publication bias, or differences in methodology.

**Graphical Abstract:** 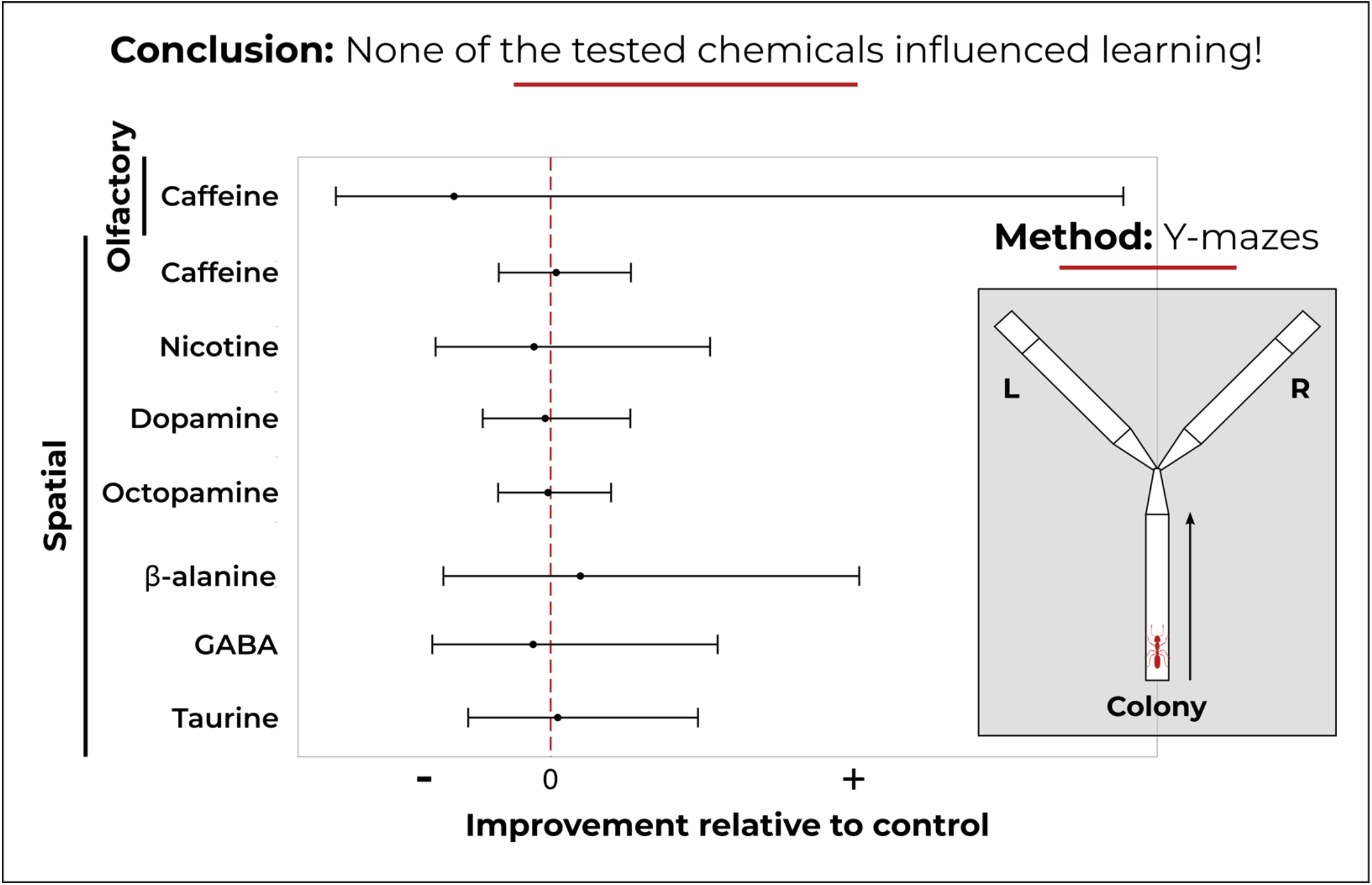

## Introduction

Plants are known to lace nectar with bioactive secondary metabolites, some of which act as neurotransmitters, binding with neuron receptor proteins, thus influencing neural activity and pollinator behaviour (Mustard, 2020). Interfering with insect neuronal signal transduction is thought to increase pollination and seed dispersal (Wink, 2018). For example, caffeine causes bees to form stronger, longer-lasting associations between odours and rewards, although such effects tend to be short-lived (Wright *et al*., 2013; Arnold *et al*., 2021). Additionally, it leads to bees overestimating resource quality, increasing foraging frequency and recruitment (Singaravelan *et al*., 2005; Couvillon *et al*., 2015; Thomson, Draguleasa & Tan, 2015).

Similarly, lacing food with β-alanine and GABA has been reported to improve associative learning and memory retention in bees. However, when ingested prior to conditioning, GABA, β-alanine and taurine hindered learning, but not memory retention, which was surprisingly improved by β-alanine and taurine (Carlesso *et al*., 2021). Dopamine and octopamine, neuromodulators in the central nervous system of invertebrates, are involved in information flow regarding food source quality, with octopamine showing an increased use of private information in bees (Linn *et al*., 2020). Octopamine and dopamine receptors have been linked to appetitive learning, with artificial increases of dopamine increasing the value of sucrose solution and improving olfactory learning and memory retrieval in both wasps and bees (Lenschow *et al*., 2018; Huang *et al*., 2022).

The effect of secondary metabolites and neurotransmitters in modulating foraging and learning in insects is currently a very active field of research. Table 1 provides examples of the effects of seven potential neuroactive chemicals on learning and memory across the Hymenoptera, whilst highlighting the significant bias towards honeybees and bumblebees as model organisms. In fact, upon extensive search, to our knowledge only six studies investigated the effects of these chemicals on ants, three of which focusing exclusively on whether the chemical elicited preference or aversion. Caffeine was shown to act as an attractant or repellent, depending on the extracts and concentrations used, likely altering food value perception (Yeoh, Dieng & Majid, 2018; Majid *et al*., 2018; Madsen & Offenberg, 2019). Furthermore, both caffeine and nicotine have been reported to improve conditioning and memory, albeit while decreasing food consumption (Cammaerts, Rachidi & Gosset, 2014a, 2014b). More recently, dopamine has been linked to long-term memory consolidation and octopamine to appetitive learning of olfactory cues (Wissink & Nehring, 2021).

**Table 1.**
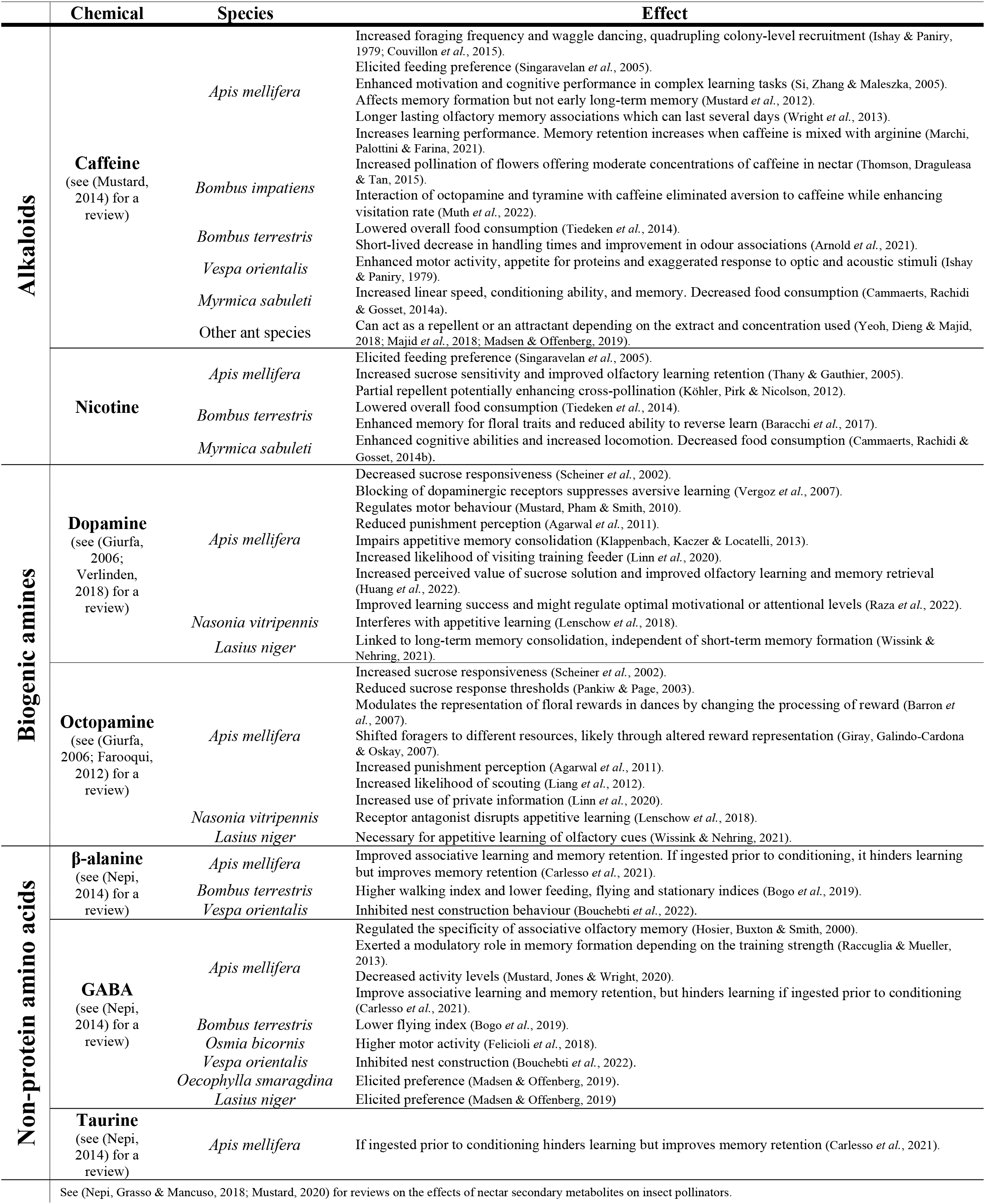
Overview of the effects of neuroactive chemicals on learning and memory in Hymenoptera.

Thus far, global invasive ant control attempts have cost over 10 billion euros (Angulo *et al*., 2022). *Linepithema humile* (Mayr, 1868) is one of the most damaging invasive alien species worldwide (Lowe *et al*., 2000), and the fourth most costly invasive ant species (Angulo *et al*., 2022). Being both ecologically and economically damaging, these ants have become a top priority for conservation programmes (Hoffmann *et al*., 2016). Nevertheless, eradication attempts have often met with failure (Souza *et al*., 2008; Hoffmann, 2011), as competition with natural food sources leads to a lack of sustained bait consumption (Rust, Reierson & Klotz, 2003; Silverman & Brightwell, 2008; Nyamukondiwa & Addison, 2011).

One way of increasing preference for, and consumption of, target foods is to target learning, a critical cognitive ability which, if exploited, can be used to steer preference (Farina *et al*., 2020). Associative learning, one of the most important types of learning, links an unconditional stimulus (any stimulus which, without learning, causes a response) with a conditional stimulus (one which can be perceived, but does not by itself result in a response). Once linked, sensing the conditional stimulus results in a similar response to the one caused by the unconditional stimulus (Pavlov, 1927; Rescorla & Wagner, 1972; Dickinson, 2012).

Ants use chemical, olfactory, and visual cues when foraging (Aron, Deneubourg & Pasteels, 1988; Roces, 1990; Czaczkes *et al*., 2014; Arenas & Roces, 2018), acquiring landmark information and building complex navigational routes (Helmy & Jander, 2003; Graham & Collett, 2006; Knaden & Graham, 2016; Wystrach *et al*., 2020). They are strong associative learners, requiring as little as one experience to form a memory which may last for up to three days (Dupuy *et al*., 2006; Josens, Eschbach & Giurfa, 2009; Huber & Knaden, 2018; Oberhauser *et al*., 2019; Piqueret, Sandoz & d’Ettorre, 2019; Czaczkes & Kumar, 2020). Specifically, *L. humile* have been shown to be incredibly fast learners, requiring as little as two experiences for 84% of the studied individuals to successfully associate a side of a Y-maze with the presence of a reward. Similar results were observed in olfactory learning, in this case with a single experience, and long-term memories were shown to last up to two days (Rossi *et al*., 2020; Wagner, Galante & Czaczkes, 2022). Enhancing or disrupting cognitive abilities, especially learning, could be a key step towards improving invasive species control.

Here, we test the effects of seven potential neuroactive chemicals (two alkaloids: caffeine and nicotine; two biogenic amines: dopamine and octopamine; three non-protein amino acids: β-alanine, GABA, and taurine) on the cognitive abilities of invasive *L. humile* in a laboratory setting. We mainly focus on short-term effects on spatial associative learning, as previous work suggests there is little room for improvement when it comes to olfactory associative learning in a laboratory setting (Wagner, Galante & Czaczkes, 2022). Improving ant navigational skills could lead to sustained bait consumption by improving both foraging and recruiting of toxicant-laced baits. The motivation for this study was potential future application in an invasive species management setting. We thus focused on effects which manifest directly after consumption, without the need for pre-treatment or topical application.

## Materials and Methods

### Colony maintenance

*Linepithema humile* (Mayr, 1868) were collected from Portugal (Proença-a-Nova and Alcácer do Sal) and Spain (Girona) between April 2021 and April 2022. Ants were split into colony fragments (henceforth colonies), containing three or more queens and 200-1000 workers, kept in non-airtight plastic boxes (32.5 × 22.2 × 11.4 cm) with a plaster of Paris floor and PTFE coated walls. 15mL red transparent plastic tubes, partly filled with water, plugged with cotton, were provided as nests. Ants were maintained on a 12:12 light:dark cycle at room temperature (21-26 °C) with *ad libitum* access to water. Between experiments, ants were fed *ad libitum* 0.5M sucrose solution and *Drosophila melanogaster* twice a week. During experiments, ants were fed once a week and deprived of carbohydrates for four to five days prior to testing, ensuring high foraging motivation. Experiments were conducted between March 2022 and September 2022 using 18 colonies divided into donor/recipient pairs. Donor colonies were kept naïve, never exposed to any of the chemicals used. During testing, focal ants left the donor colony, but returned to the recipient colony, where they unloaded the contents of their crop.

### Chemicals and solutions

Caffeine (CAS 58-08-2), nicotine (CAS 65-30-5), dopamine (CAS 62-31-7), octopamine (CAS 770-05-8), β-alanine (CAS 107-95-9), GABA (CAS 56-12-2), taurine (CAS 107-35-7) and ascorbic acid (CAS 50-81-7) were obtained from Sigma-Aldrich (Taufkirchen, Germany). 1M sucrose solutions (Südzucker AG, Mannheim, Germany) mixed with a single chemical were used as treatments. Identical 1M sucrose solutions were used as controls across all experiments. Chemical concentrations were chosen based on previous reports of their effects on Hymenopterans. Caffeine has shown neuroactive effects at a wide range of concentrations (Mustard, 2014). Therefore, 1.29μmol ml^−1^, a moderately high concentration, ten-fold the naturally occurring one, was used (Singaravelan *et al*., 2005; Mustard *et al*., 2012). Nicotine was used at 0.02μmol ml^−1^ (Thany & Gauthier, 2005; Cammaerts, Rachidi & Gosset, 2014b; Baracchi *et al*., 2017). 10.55μmol ml^−1^ of dopamine or octopamine were mixed with 9.94μmol ml^−1^ of ascorbic acid to reduce oxidation of the biogenic amines (Scheiner *et al*., 2002; Linn *et al*., 2020). β-alanine, GABA and taurine were used at 0.27μmol ml^−1^, 0.73μmol ml^−1^ and 0.32μmol ml^−1^, respectively (Carlesso *et al*., 2021). A double-blind procedure was applied to all solutions used to minimize experimenter bias.

### Y-maze experimental setup – Spatial learning

Y-mazes (three 10cm long, 1cm wide arms, tapering to 2mm at the bifurcation) were used to assess the effects of each chemical on spatial memory and learning (Czaczkes, 2018). Each donor colony was connected to a Y-maze via a drawbridge, both covered in unscented disposable paper overlays. A drop of sucrose solution (positive stimulus), either the control or the treatment, was placed at the end of one of the maze arms, and a drop of water (neutral stimulus) on the opposing arm. The first two ants willing to walk up the drawbridge were allowed onto the Y-maze and marked with acrylic paint while drinking the sucrose solution. Upon satiation, ants were not allowed back into their original donor colony. Rather, they were allowed to return to the paired recipient colony, where they offloaded the content of their crop. Meanwhile, the Y-maze paper overlays were replaced, to remove any pheromone trails left behind, and the solution drops reapplied to their original maze arm. Following trophallaxis, within 0-30 minutes since the end of the first visit, one of the two marked ants was allowed back onto the Y-maze. Its initial decision was recorded as the first maze arm in which it crossed a 2cm reference line, and its final decision as the maze arm containing the drop it first touched. To account for a potential time-dependent effect of the neuroactive chemicals tested, the second marked ant was only allowed back onto the Y-maze 31-60 minutes after the end of the first visit. For the caffeine experiment, instead of two, five ants were initially marked. In this case, each ant’s second visit occurred in increasing 30-minute intervals going up to over 120 minutes since the end of its first visit. This followed previous literature reporting delayed caffeine effects ranging between 30-120 minutes in honey bees (Mustard *et al*., 2012; Gong *et al*., 2021). From the second visit onwards, ants were allowed back onto the Y-maze as soon as possible. In total, each ant carried out five consecutive visits to the Y-maze: an initial one where it was marked and no data was collected, and four others where their choice was recorded. The treatment used, the Y-maze arm in which it was located and the elapsed time since the end of the first visit were randomly assigned to each individual following a full factorial design. A total of 481 individuals were tested across seven experiments.

### Y-maze experimental setup – Olfactory learning

Y-mazes were also used to study the effects of caffeine on olfactory memory and learning. Scented paper overlays, used during testing, were stored in airtight plastic boxes (19.4 × 13.8 × 6.6 cm) containing an open glass petri-dish with 0.5mL of either strawberry or apple food flavouring (Seeger, Springe, Germany) for at least a week prior to use. An individual ant from a donor colony was allowed onto a 10cm linear runway covered by a scented paper overlay offering a sucrose solution drop (positive stimulus), either pure or laced with 1.29μmol ml^−1^ caffeine, at the end. The ant was marked while drinking and, upon satiation, was allowed to return to the paired recipient colony to offload its crop content. After unloading, the marked ant was allowed onto a Y-maze offering on one arm a paper overlay scented to match the odour experienced during training, and on the other arm the opposing odour (novel stimulus). The ants’ initial and final choice was recorded as the first maze arm in which it crossed a 2cm and an 8cm reference line, respectively. The treatment used, the Y-maze arm in which it was located, and the odour associated with the reward were randomly assigned to each individual following a full factorial design, testing 96 individuals.

### Statistical analysis

All graphics and statistical analysis were generated using R version 4.2.1 (R Core Team, 2022), using the packages reshape2 (Wickham, 2007) and ggplot2 (Wickham, 2016). Analysis was conducted by multi-model inference following an information theory approach (Anderson, 2008). Generalised linear mixed models were fit using the lme4 (Bates *et al*., 2015) package with binomial error distributions and estimated marginal means and contrasts were obtained using the emmeans package (Lenth, 2022) with Bonferroni adjusted values accounting for multiple testing. An *a priori* set of hypotheses, and matching candidate models, was developed for each experiment (Table 2). The DHARMa (Hartig, 2022) package was used to inspect the global model in each set, from which all other models can be derived, assessing model fit and ensuring model assumptions were met (Burnham & Anderson, 2002). Conditional coefficients of determination, a measure of goodness of fit, were calculated for each model using the MuMIn (Bartoń, 2022) package. The AICcmodavg (Mazerolle, 2020) package was used to calculate Akaike’s information criterion, adjusted for small sample sizes (AICc), and Akaike weights (w_i_) for each model. Model-averaged parameter estimates, standard errors and confidence intervals were then computed as a weighted mean of the set of candidate models. We avoid the use of p-values, instead reporting effect size estimates and their respective 95% confidence intervals (Greenland *et al*., 2016) shown throughout the results section as (estimate [lower limit, upper limit], N = sample size).

**Table 2.** Candidate model set and corresponding *a priori* hypothesis used for multimodel inference. All models used the proportion of ants choosing the rewarded side of the Y-maze as their final decision as the response variable and included data collection date, colony identity and ant identity as random effects. Additionally, spatial learning models included experimenter and colony starvation period as random effects.

|  | Model | Biological Hypothesis |
| --- | --- | --- |
| Spatial | Null | Ants randomly choose a Y-maze arm. |
|  | Visit | Learning improves over consecutive visits. |
|  | Treatment | The neuroactive chemical interferes with learning. |
|  | Reward Side | Ants have an intrinsic predisposition towards turning left or right. |
|  | Elapsed Time | Recall strength, and therefore learning, is affected by the time memories had to consolidate. |
|  | Treatment * Elapsed Time | The effects of the neuroactive chemical on learning are time dependent. |
|  | Treatment * Visit | The neuroactive chemical interference varies with learning strength. |
|  | Maximal | All the variables of interest contribute towards learning. |
| Olfactory | Null | Ants randomly choose a Y-maze arm. |
|  | Treatment | The neuroactive chemical interferes with learning. |
|  | Reward Side | Ants have an intrinsic predisposition towards turning left or right. |
|  | Odour | Ants have an innate preference towards specific odours. |
|  | Treatment * Odour | The neuroactive chemical might affect the ant's perception of the odour. |
|  | Maximal | All the variables of interest contribute towards learning. |

## Results

The complete statistical analysis output for all experiments, and the entire dataset on which this analysis is based, is available from Zenodo (https://doi.org/10.5281/zenodo.7268444).

Binomial generalised linear models were used to check for differences between the proportion of ants choosing the rewarded side of the Y-maze as their initial versus their final decision for each experiment (conditional R^2^ range of 20-69%, N = 8). Post-hoc estimated marginal means, with Bonferroni-adjusted significance levels, based on each spatial learning model revealed small differences between initial and final decision (0.3-7.1%, N = 7). However, the same method applied to the olfactory learning experiment revealed a relatively large difference between initial (76.5% [47.2%, 92.2%], N = 96) and final decision (93.1% [69.7%, 98.8%], N = 96). Across experiments, the proportion of ants choosing the rewarded side of the maze as their final decision was always higher than that of ants doing so as their initial decision. This suggests that ants often realised that they had entered the unrewarded maze arm and corrected their decision. Such corrections imply ants recall the location of the reward and are likely learning. As our aim was to explore the effects of different neuroactive chemicals on learning all statistical analysis used final decision as the response variable.

### Ants learn to associate a reward with a scent and with a side of a Y-maze

All candidate models (Table 2) were fit using generalised linear models with binomial error distributions for each experiment. The conditional R^2^, a measure of goodness of fit, for the model which explains the most variance in the data for each set of candidate models is reported in Figure 2 (conditional R^2^ range of 12-43%, N = 8). Estimated marginal means, with Bonferroni-adjusted significance levels, averaged over the treatments used and the side of the maze in which the reward was located, show that ants can associate both the apple (80.0% [63.0%, 90.4%], N = 48) and strawberry (78.4% [63.0%, 88.5%], N = 48) scents with the presence of a sucrose reward after a single training visit. Similarly, Figure 1 shows that ants can associate the presence of a reward with a side of a Y-maze and that learning tends to increase over consecutive visits. It is worth noting that for the octopamine experiment, ants had a significant innate side bias towards turning left. This same trend was seen across all experiments, although for all others it was not statistically significant (see ESM1).

**Figure 1.**
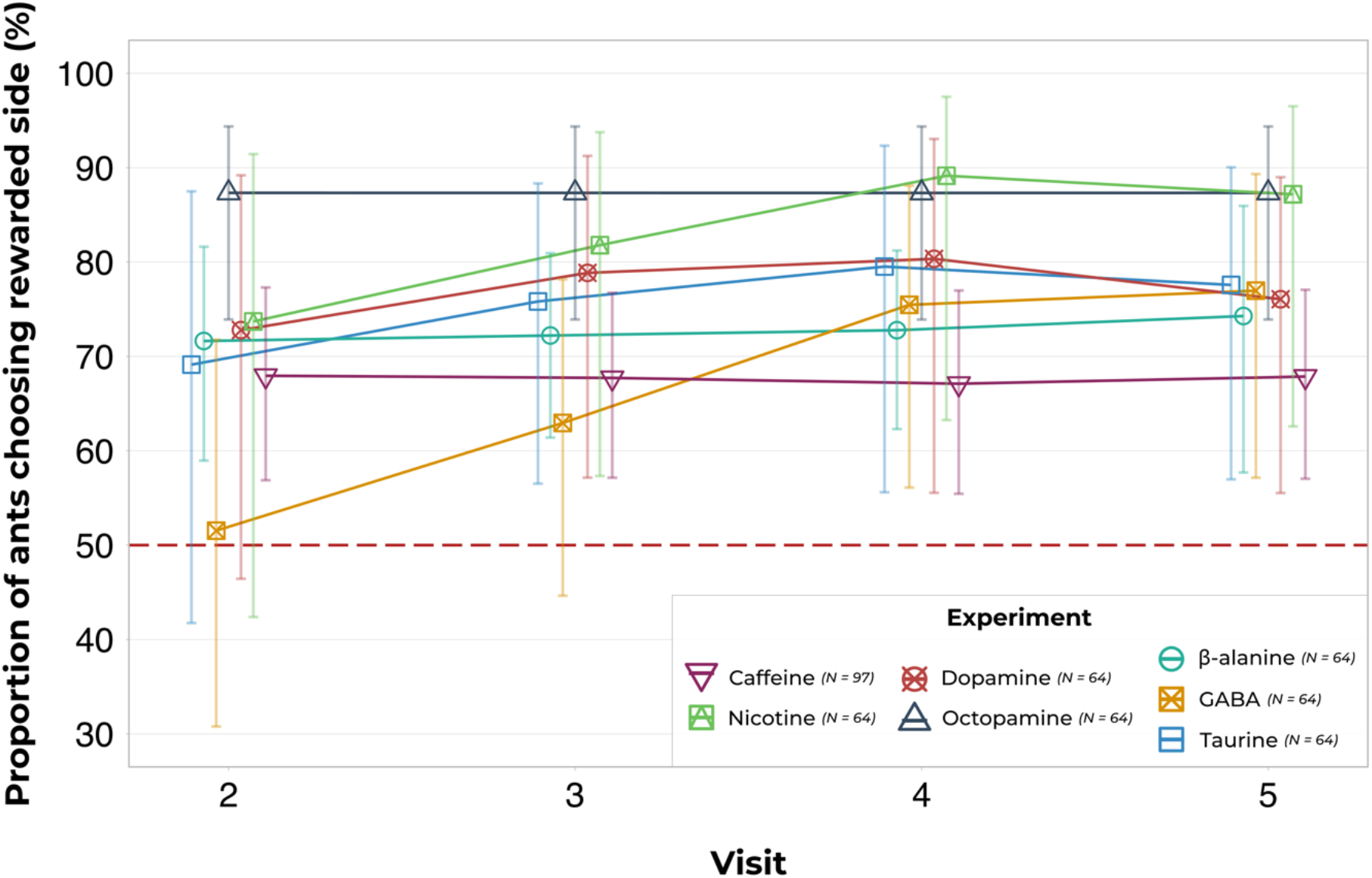
Ants learn to associate a sucrose reward with an arm of a Y-maze over consecutive visits. Shapes represent the proportion of ants (control + treatment) choosing the rewarded side of the maze as their final choice and whiskers the respective 95% unconditional confidence intervals. Estimates for each experiment were obtained from model-averaging with shrinkage and estimated marginal means were averaged over the treatments used, the side of the maze in which the reward was located and the elapsed time since the end of the first visit. This was done as the models show no significant difference between treatment, reward side or elapsed time. The exception to this being the octopamine experiment which showed a significant side bias towards the left (L = 93%, R = 77%). However, since even ants with the reward on the right were able to learn the association, we average both sides. If the confidence intervals of each estimate include 50% (red dashed horizontal line), ants are considered to choose an arm of the Y-maze at random and therefore likely did not learn. Significance levels were adjusted using Bonferroni correction for multiple testing.

### None of the chemicals tested influenced learning

Parameter estimates for each experiment were obtained from model-averaging with shrinkage as odds ratios. Odds are the probability of an event occurring divided by the probability of the event not occurring. Odds ratios compare two odds, testing how the relationship between these two odds change given different conditions. Figure 2 shows the estimated odds ratios for each experiment comparing the odds of an ant under the influence of each chemical choosing the rewarded side of the Y-maze against the odds of an ant under the influence of the respective control treatment doing so, if all other variables are kept constant. Odds ratios of 1 indicate no difference between the treatment and its control, whilst odds ratios > 1 or < 1 indicate that ants are more or less likely, respectively, to choose the rewarded side of the Y-maze under the influence of the neuroactive chemical when compared to the control. Our results suggest that none of the chemicals used interferes with *L. humile* associative learning.

**Figure 2.**
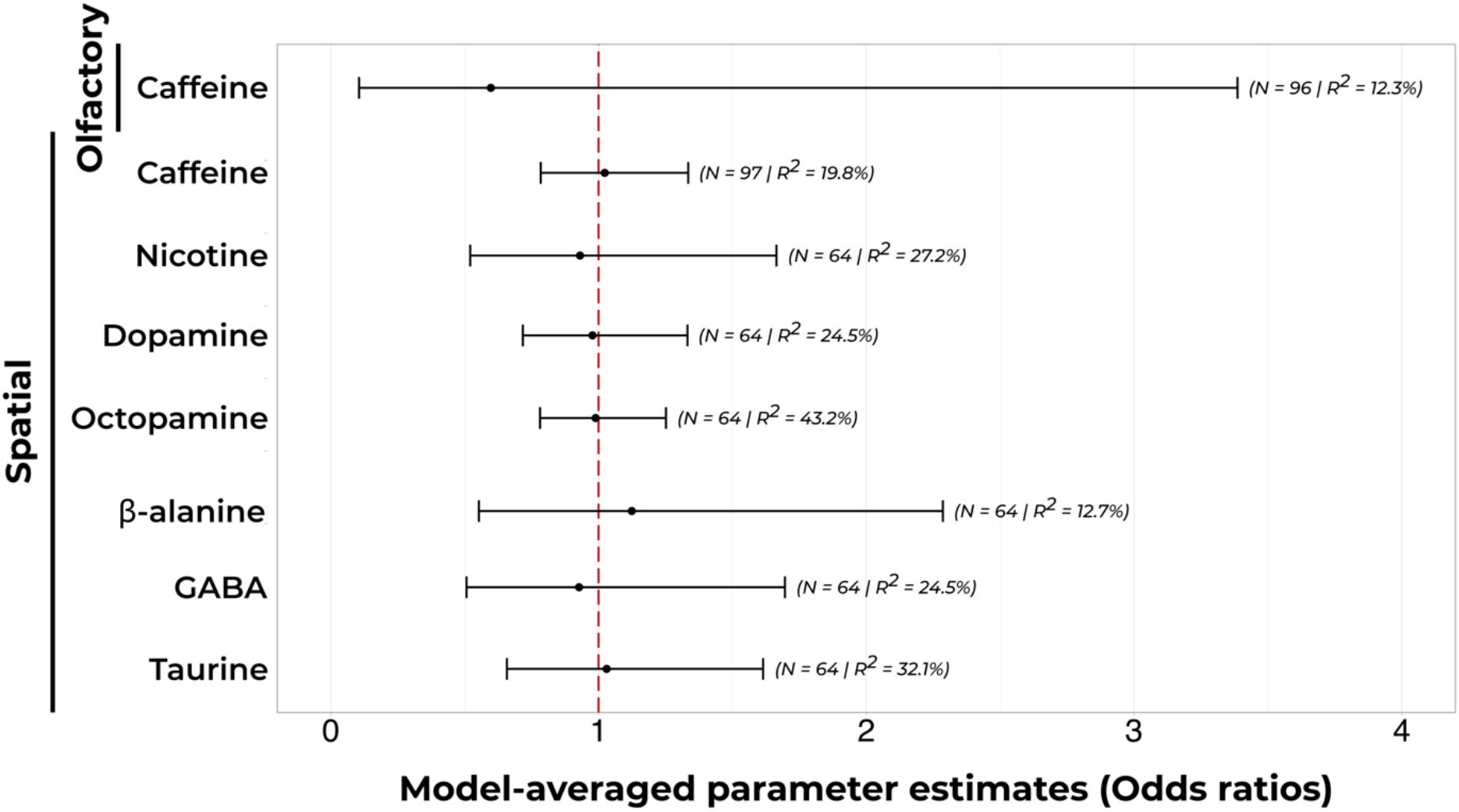
Effect of seven potential neuroactive chemicals on the olfactory and spatial associative learning of *L. humile*. Circles represent the estimates obtained from model-averaging with shrinkage and whiskers the 95% unconditional confidence intervals. The odds ratio compares the odds (probability of an event occurring divided by the probability of the event not occurring) of the ants choosing the rewarded side of the Y-maze under the influence of each neuroactive chemical against those of the corresponding control treatment. Odds ratios of 1 (red dashed vertical line) indicate no difference between the treatment and its control, whilst odds ratios > 1 or < 1 indicate that ants are more or less likely, respectively, to choose the rewarded side of the Y-maze under the influence of the neuroactive chemical when compared to the control. If the 95% confidence intervals include an odds ratio of 1 there is no significant difference between treatment and control. R^2^ refers to the goodness of fit of the model which explains the most variance in the data for each set of candidate models.

Additionally, we collected data regarding the time each ant took from entering the Y-maze until it reached the reward (“In Duration”) and the elapsed time since each ant finished drinking the reward until it reached the entrance of the maze (“Out Duration”). Since the Y-maze represents a relatively short and straightforward distance, it is hard to detect small variations between treatments. Nevertheless, we performed a simple survival analysis, computing the probability of each ant reaching the reward or the nest, at specific points in time. Ants treated with β-alanine returned to the nest faster than control treated ants. However, although statistically significant, this effect is relatively small, but suggests β-alanine is a promising neuroactive chemical for further testing. Similarly, it is worth noting that dopamine and octopamine seem to increase the time ants take to reach the reward, although this effect was not statistically significant (see ESM1 for detailed analysis and figures).

## Discussion

*L. humile* are incredibly effective associative learners (Rossi *et al*., 2020; Wagner, Galante & Czaczkes, 2022). Here, we show that a single training visit to a Y-maze is often enough for ants to develop a spatial association between the presence of a reward and an arm of the maze. Ants often correct their initial decision, further suggesting they are in fact learning. Over consecutive visits, the proportion of ants choosing the rewarded side of the maze increases until it plateaus, with three and four training visits showing similarly strong learning. Furthermore, after a single training visit, *L. humile* show an extremely strong preference for the scent they were trained with over a novel one. These results support previous work suggesting ants require as little as one experience to form a memory retaining it for up to three days (Dupuy *et al*., 2006; Josens, Eschbach & Giurfa, 2009; Huber & Knaden, 2018; Oberhauser *et al*., 2019; Piqueret, Sandoz & d’Ettorre, 2019; Czaczkes & Kumar, 2020). Furthermore, across all experiments, ants seem to have an innate preference towards turning left, even if in most cases this does not hinder learning. Such preference is likely linked to brain lateralisation with a preference towards the left being shown in ants previously (Hunt *et al*., 2014).

None of the seven potential neuroactive chemicals tested showed a significant effect on spatial learning, with caffeine also not influencing olfactory associative learning. This is in contrast to the extensive literature on the effects of these chemicals on Hymenopterans (Table 1). Honeybees prefer sucrose solutions laced with up to 0.52μmol ml^−1^ caffeine (Singaravelan *et al*., 2005) with topically delivered caffeine improving both motivation and cognitive performance of complex learning tasks at vastly greater concentrations (Si, Zhang & Maleszka, 2005). Similarly, 5.15μmol ml^−1^ caffeine was reported to increase conditioning ability and memory in ants (Cammaerts, Rachidi & Gosset, 2014a). However, due to a positive publication bias (Nissen *et al*., 2016; Mlinarić, Horvat & Smolčić, 2017), it is extremely hard to find null results to contextualise our findings. As an example, two unpublished Master’s theses have studied the chronic effects of caffeine on honeybee learning, and both suggest a general lack of effect on learning performance (Malechuk, 2009; Yusaf, 2012).

The lack of effect we found in this study does not rule out these chemicals as influencing spatial learning in ants (see Box 1). Although the chemical concentrations used were chosen based on previous literature showing their effects on Hymenoptera, it could be that we missed the concentration at which they influence learning and memory. For instance, unnaturally high concentrations of nicotine deterred bumblebees, but lower nectar-relevant concentrations lead to attraction (Baracchi *et al*., 2017). Furthermore, it is likely that the effects of the neuroactive chemicals used are time-dependent, and therefore this study could have missed the chemical activation window. In fact, honeybees fed 1.04μmol ml^−1^ caffeine were more likely to remember a conditioned scent than the respective control at both 24 and 72 hours after conditioning, but not 10 minutes after conditioning (Wright *et al*., 2013). Nevertheless, honeybees fed 0.05μmol ml^−1^ and 0.51μmol ml^−1^ caffeine showed stronger memory retention at two and 24 hours post-treatment, with more recent treatment resulting in stronger recall (Gong *et al*., 2021). Similarly, high concentrations of caffeine (> 10.32μmol ml^−1^) lead to a significant decrease in memory retention five minutes post-treatment (Mustard *et al*., 2012). Here, we specifically focussed on short time frames, as we were exploring the potential for neuroactive chemicals to improve bait consumption in the field. Baiting is, however, both costly and time sensitive, with modern hydrogel bead delivery systems desiccating quickly, lasting up to two hours (Cabrera *et al*., 2021). For this reason, we focused on short-term effects with an activation window of up to two hours.

**Box 1** — Future directions

Neuroactive chemicals are likely to influence learning and memory in ants. However, our work suggests that such effects might not manifest over short time periods. Thus, steering ant preference with neuroactive chemicals might not be ideally suited to application in pest control. Nevertheless, understanding how these chemicals influence learning and memory still offers significant mechanistic insights into the insect brain. Here, we propose some potential avenues of exploration which we think would be of particular interest:

- Focusing on olfactory learning, which is thought to take place in the acetylcholine receptor-rich antennal lobes and mushroom bodies.
- Using lower sucrose concentrations would reduce motivation, in theory decreasing learning speed or quality, which could help studying subtle effects induced by the chemicals – especially in the face of ceiling effects caused by excellent olfactory learning.
- Using different, more complex tasks, such as reversal learning or navigation in an open field (Galante et al. In prep.) would require more neural pathways to be activated and therefore could help expose effects induced by the chemicals.
- Testing learning, but also its extinction, could provide insights into how these chemicals impact long-term memory formation, consolidation, and retention.
- Using different concentrations and combinations of various nectar secondary metabolites seems to be promising – for example combining caffeine with arginine or octopamine and tyramine (Marchi, Palottini & Farina, 2021; Muth et al., 2022).
- β-alanine is a promising chemical for further testing, as it caused a small but significant reduction in return time to the nest.

There is a significant literature bias towards bees as model organisms, often using the proboscis extension response (PER) paradigm and focusing on olfactory associative learning. It could thus be that the contrast between our results and much of the published literature stems from species specific differences and/or methodological ones. It is possible that the chemicals studied target specific neurological pathways that are activated during PER experiments, but not during the ones we conducted. A wide range of acute doses of caffeine has been shown to affect learning but not memory in honeybees (Mustard *et al*., 2012). This suggests neuroactive chemicals have high specificity and therefore it is likely that different tasks are disrupted differently. In fact, caffeine and nicotine target acetylcholine receptors (AChR) which are abundant in the antennal lobes and mushroom bodies, the same areas thought to be responsible for appetitive olfactory learning in bees (MaBouDi *et al*., 2017; Mustard, 2020). Contrastingly, spatial learning is thought to mainly occur at the level of the central complex (Ofstad, Zuker & Reiser, 2011), which might have less AChR expressed and might therefore remain unaffected by chemicals that target it. The lack of an effect on olfactory learning in the current study could also be due to a ceiling effect: we replicate the previous finding that there is little to no room for improvement when it comes to olfactory associative learning in *L. humile* ants (Wagner, Galante & Czaczkes, 2022). Thus, even if caffeine does improve olfactory learning in these ants, it would be hard for such an effect to be visible due to their already excellent natural learning.

Finally, recent studies suggest that, like in plants, a combination of different neuroactive chemicals might be key towards manipulating behaviour. Honeybees fed 0.05μmol ml^−1^ or 0.16μmol ml^−1^ of caffeine showed improved learning performance, but no change in memory retention unless caffeine was mixed with arginine (Marchi, Palottini & Farina, 2021). Moreover, octopamine and tyramine mixed with caffeine altered bumblebee behaviour, but not when present individually (Muth *et al*., 2022). However, considering the infinite possible combinations of chemicals at different concentrations, it seems that using neuroactive chemicals to artificially manipulate ant behaviour might not be straightforward. Nevertheless, many promising avenues of research remain unexplored (see Box 1). Understanding how neuroactive chemicals influence learning and memory still offers significant mechanistic insights which could be leveraged towards improving invasive ant control.

## Acknowledgements

We thank E. Sequeira and S. Abril for ant collection, A. Koch, L. Neubauer and S. Kau for help with data collection, and D. Baracchi and M. De Agrò for manuscript feedback. H. G was supported by an ERC Starting Grant to T.J.C (H2020-EU.1.1. #948181). T. J. C. was supported by a Heisenberg Fellowship from the Deutsche Forschungsgemeinschaft (CZ 237 / 4-1).

